# Cross-family transfer of the Arabidopsis cell-surface immune receptor LORE to tomato confers sensing of 3-hydroxylated fatty acids and enhanced disease resistance

**DOI:** 10.1101/2024.04.19.590144

**Authors:** Sabine Eschrig, Parvinderdeep S. Kahlon, Carlos Agius, Andrea Holzer, Ralph Hückelhoven, Claus Schwechheimer, Stefanie Ranf

**Affiliations:** Technical University of Munich, TUM School of Life Sciences, Chair of Phytopathology, Freising-Weihenstephan, Germany; Technical University of Munich, TUM School of Life Sciences, Chair of Plant Systems Biology, Freising-Weihenstephan, Germany; University of Fribourg, Department of Biology, Fribourg, Switzerland; Current affiliation: Wageningen University and Research, Laboratory of Plant Physiology, Plant Sciences Group, Wageningen, The Netherlands

**Keywords:** Cross-family PRR transfer, resistance engineering, pattern recognition receptor, crop disease resistance, plant immunity

## Abstract

Plant pathogens pose a high risk of yield losses and threaten food security. Technological and scientific advances have improved our understanding of the molecular processes underlying host-pathogen interactions, which paves the way for new strategies in crop disease management beyond the limits of conventional breeding. Cross-family transfer of immune receptor genes is one such strategy that takes advantage of common plant immune signaling pathways to improve disease resistance in crops. Sensing of microbe- or host damage-associated molecular patterns (MAMPs/DAMPs) by plasma membrane-resident pattern recognition receptors (PRR) activates pattern-triggered immunity (PTI) and restricts the spread of a broad spectrum of pathogens in the host plant. In the model plant *Arabidopsis thaliana*, the S-domain receptor-like kinase LIPOOLIGOSACCHARIDE-SPECIFIC REDUCED ELICITATION (*At*LORE, SD1-29) functions as PRR, which senses medium chain-length 3-hydroxylated fatty acids (mc-3-OH-FAs), such as 3-OH-C10:0, and 3-hydroxyalkanoates (HAAs) of microbial origin to activate PTI. In this study, we show that ectopic expression of the Brassicaceae-specific PRR *At*LORE in the solanaceous crop species *Solanum lycopersicum* cv. M82 leads to the gain of 3-OH-C10:0 immune sensing without altering plant development. *AtLORE*-transgenic tomato shows enhanced resistance against *Pseudomonas syringae* pv. *tomato* DC3000 and *Alternaria solani* NL03003. Applying 3-OH-C10:0 to the soil before infection induces resistance against the oomycete pathogen *Phytophthora infestans* Pi100 and further enhances resistance to *A. solani* NL03003. Our study proposes a potential application of *AtLORE*-transgenic crop plants and mc-3-OH-FAs as resistance-inducing bio-stimulants in disease management.

## INTRODUCTION

Since the early days of agriculture, plant diseases have affected food production and security for humankind. Despite substantial advances in agricultural practices, global crop production still suffers significant economic losses due to plant diseases, with the highest losses in parts of the world where food security is already at risk (Oerke, 2005; Savary *et al*., 2019). Historic disease outbreaks illustrate the extent of devastation plant pathogens can cause (van Esse *et al*., 2020). The Irish potato famine caused by the oomycete *Phytophthora infestans* (Yoshida *et al*., 2013) or the great Bengal famine caused by the fungal Brown spot disease of rice (Padmanabhan, 1973; Surendhar *et al*., 2021) claimed millions of lives and led to mass emigrations. Fungal Fusarium wilt disease almost wiped out banana cultivation (Pegg *et al*., 2019), while the papaya industry in Hawaii was massively threatened by infection with the ringspot virus (Gonsalves, 1998).

Tomato is one of the most important and widely consumed vegetable crops worldwide and is susceptible to several plant pathogens (Anders *et al*., 2021; Panno *et al*., 2021; Bozbuga *et al*., 2022). Bacterial diseases affecting tomato cultivation include bacterial speck caused by *Pseudomonas syringae* pv. *tomato* (*Pst*), bacterial spot caused by *Xanthomonas* species or bacterial wilt caused by *Ralstonia solanacearum* (Wang *et al*., 2018; Panno *et al*., 2021). The oomycete *Phytophthora infestans* (late blight) and the fungus *Alternaria solani* (early blight) are widespread in tomato cultivation and contribute to significant yield losses if not controlled (Adhikari *et al*., 2017; Mazumdar *et al*., 2021).

The basis for the effective and sustainable control of plant diseases today and in the future is a detailed molecular understanding of the interactions between plants and pathogens. The last 30 years have seen immense advances in research investigating the plant immune system (Ngou *et al*., 2022). Plants evolved an effective, genetically determined immune system consisting of preformed barriers and induced defense mechanisms (Dangl & Jones, 2001; Jones & Dangl, 2006; Ngou *et al*., 2022). Pattern-triggered immunity (PTI) is based on the recognition of conserved microbe-or damage-associated epitopes and phytocytokines by pattern recognition receptors (PRRs) (Ngou *et al*., 2022). PRR activation induces a broad-spectrum defense response via transcriptional, metabolic and hormonal reprogramming, both locally at the site of infection and systemically in distal tissues (Tsuda & Somssich, 2015; Vlot *et al*., 2021; Ngou *et al*., 2022). Adapted pathogens secrete effector molecules into their hosts to manipulate PTI and host physiology to their advantage. Detection of these microbial manipulations by plant resistance (R) proteins activates effector-triggered immunity (ETI) (Ngou *et al*., 2022). PTI and ETI are strongly intertwined and cooperatively contribute to disease resistance (Ngou *et al*., 2021; Yuan *et al*., 2021b). The central role of PTI is underscored by the fact that PTI is essential for a full ETI response, which is accomplished largely through the potentiation of PTI (Ngou *et al*., 2021; Tian *et al*., 2021; Yuan *et al*., 2021b). Basic molecular research in the field of plant immunity improves our understanding of the complex interaction between plants and pathogens and paves the way for new strategies to combat crop diseases.

Effective crop protection combines cultural, chemical, biological and genetic control methods (Sharma *et al*., 2022). While chemical control has become more challenging due to increasing pathogen resistance to pesticides and the rise of concerns regarding their potential toxicity, environmental impact and human health risks, genetic crop protection offers a more environmentally friendly and less labor and cost-intensive strategy (Dangl *et al*., 2013; Isman, 2019; van Esse *et al*., 2020). Historically, genetic host resistance has been achieved by the identification of new quantitative trait loci (QTL) for basal resistance or *R* genes in the gene pools of wild relatives and their introgression into elite crop cultivars. However, lack of sexual compatibility, long generation times, and difficult introgression into polyploid species limit the breeding processes (Dangl *et al*., 2013). Moreover, since race-specific *R* genes drive the adaptation of pathogen populations to overcome resistance, *R* gene-based breeding strategies often lead to short-lived resistance in the field (Dangl *et al*., 2013; Huang & Zimmerli, 2014). Besides advanced marker-assisted resistance breeding, genome editing and transgenic approaches became a newly emerging field in genetic crop protection (Pickett, 2016; Wang *et al*., 2019). They can effectively enhance disease resistance by the addition or modulation of defense components and have shorter development times compared to conventional breeding techniques (Gurr & Rushton, 2005; van Esse *et al*., 2020; Sharma *et al*., 2022). Although the application of genetically modified organisms is subject to strict legal regulations and public acceptance across the globe varies and can be locally low, proof-of-concept studies demonstrate the efficacy of such genetic approaches and their relevance for future crop protection measures (Pickett, 2016; Wang *et al*., 2019; Anders *et al*., 2021; Bubolz *et al*., 2022). For instance, genome editing of the resistance gene *mlo* in tomato provided resistance to powdery mildew (Nekrasov *et al*., 2017). Transgenic ‘Rainbow’ papaya is a commercially available ringspot virus-resistant variety overexpressing a viral coat protein, which saved the papaya industry in Hawaii in the late 1990s (Fitch *et al*., 1992; Ferreira *et al*., 2002). Inter-species transfer of the resistance gene *Bs2* from pepper to tomato improved resistance against bacterial spot caused by *Xanthomonas* spp. (Horvath *et al*., 2012).

As PTI is critical for plant health (Yuan *et al*., 2021b) and recognized elicitors are considered less likely to adopt mutations to evade PRR recognition (Huang & Zimmerli, 2014), PTI-based crop engineering is a great alternative to *R* gene transfer and may lead to broad-spectrum and, hence, more durable resistance. PRRs are attractive targets for genetic resistance engineering, as they can be transferred into host plants to expand their receptor repertoire (Huang & Zimmerli, 2014; Ranf, 2018). The great potential of cross-family PRR transfer was first demonstrated by the integration of the Brassicaceae-specific PRR EF-Tu RECEPTOR (EFR) from *Arabidopsis thaliana* into *Nicotiana benthamiana* and tomato, conferring increased disease resistance against various bacterial pathogens (Lacombe *et al*., 2010). *At*EFR was later integrated into rice, wheat, potato, orange and apple (Lacombe *et al*., 2010; Lu *et al*., 2015; Schoonbeek *et al*., 2015; Schwessinger *et al*., 2015; Boschi *et al*., 2017; Mitre *et al*., 2021; Piazza *et al*., 2021). The effectiveness of *At*EFR-transgenic tomato against bacterial wilt disease caused by *Ralstonia solanacearum* and bacterial spot caused by *Xanthomonas perforans* was confirmed in field trials in the US (Kunwar *et al*., 2018). Another example is the transfer of the rice immune receptor kinase Xa21 to orange, tomato and banana to enhance resistance against wilt diseases (Mendes *et al*., 2010; Afroz *et al*., 2011; Tripathi *et al*., 2014). Expression of the extracellular ATP (eATP) receptor DOES NOT RESPOND TO NUCLEOTIDES 1 (DORN1 or LecRK-1.9) of *A. thaliana* in *Solanum tuberosum* and *N. benthamiana* increased resistance to *P. infestans* (Bouwmeester *et al*., 2014; Wang *et al*., 2016). Finally, citrus species expressing the FLS2 receptor from *N. benthamiana* are more resistant to citrus canker caused by *Xanthomonas citri* (Hao *et al*., 2016).

In the model plant *A. thaliana*, we previously identified the S-domain receptor-like kinase (SD-RLK) LIPOOLIGOSACCHARIDE-SPECIFIC REDUCED ELICITATION (*At*LORE, SD1-29, AT1G61380) as a PRR (Ranf *et al*., 2015) that senses bacterial 3-hydroxylated fatty acids (mc-3-OH-FAs) and 3-hydroxyalkanoates (HAAs) of medium chain-length ranging from C8 to C12 (Kutschera *et al*., 2019; Schellenberger *et al*., 2021). Free 3-hydroxydecanoic acid (3-OH-C10:0) is the strongest elicitor of *At*LORE-dependent immunity (Kutschera *et al*., 2019). Mc-3-OH-FAs bound to acyl carrier protein (ACP) and coenzyme A (CoA) are general intermediates of fatty acid metabolism and serve as building blocks for various lipidic microbial compounds, which potentially release free mc-3-OH-FAs (Kutschera *et al*., 2019; Cho *et al*., 2020). *At*LORE homomerizes (Eschrig *et al*., 2024), and upon ligand sensing, activates the activates the receptor-like cytoplasmic kinases PBS1-LIKE 34/35/36 (*At*PBL34/35/36) and RPM1-INDUCED PROTEIN KINASE (*At*RIPK) for downstream signaling and the LORE-ASSOCIATED PROTEIN PHOSPHATASE (*At*LOPP) for signaling regulation (Luo *et al*., 2020; Li *et al*., 2021; Wang *et al*., 2023). *At*LORE promotes resistance against *Pst* DC3000 and *R. solanacearum* in *A. thaliana* (Kutschera *et al*., 2019; Wang *et al*., 2023). *Pst* DC3000 resistance can be further enhanced by pretreatment with 3-OH-C10:0 or HAAs before infection (Kutschera *et al*., 2019; Schellenberger *et al*., 2021).

*LORE* is phylogenetically restricted to Brassicaceae (Ranf *et al*., 2015; Eschrig *et al*., 2024) and thus an excellent model for application in genetic crop disease management via cross-family gene transfer. We have previously shown that transient expression of *At*LORE in solanaceous *Nicotiana benthamiana* confers sensitivity to mc-3-OH-FAs (Ranf *et al*., 2015; Eschrig *et al*., 2024). In this study, we generated transgenic *AtLORE*-overexpressing *S. lycopersicum* cv. M82 to investigate the feasibility of functionally transferring an SD-type PRR across plant families into an agriculturally relevant crop. We tested whether *AtLORE*-transgenic tomato is more resistant to the adapted tomato pathogen *Pst*, a prominent producer of 3-OH-C10:0. Furthermore, we investigated whether soil application of 3-OH-C10:0 induces systemic resistance of *AtLORE*-transgenic tomato to other tomato pathogens, which might not directly activate LORE-dependent immunity, such as the fungus *A. solani* or the oomycete *P. infestans*.

## RESULTS

### Generation of stable transgenic *S. lycopersicum* overexpressing *AtLORE*

The finding that LORE confers resistance to *P. syringae* and *R. solanacearum* in *A. thaliana* (Kutschera *et al*., 2019; Wang *et al*., 2023) and that *At*LORE is functional in *N. benthamiana* upon transient expression (Ranf *et al*., 2015; Eschrig *et al*., 2024) raised the question of whether stable expression of *AtLORE* in *S. lycopersicum* also confers gain of mc-3-OH-FA immune sensing and increased resistance to tomato pathogens. Therefore, we generated transgenic *S. lycopersicum* cv. M82 via Agrobacteria-mediated transformation, callus culture and regeneration of transgenic plants. We ectopically overexpressed *AtLORE* under control of the strong Cauliflower Mosaic Virus 35S (CaMV 35S) promoter (*At*LORE-OE lines) or integrated the T-DNA from an empty vector (EV lines) as negative control (**Fig 1A**). Four regenerated *At*LORE-OE plants from different calli and one EV plant were grown until fruiting in the greenhouse and seeds were harvested from ripe tomato fruits (**Fig S1**). No obvious visible differences in growth or fruiting behavior were observed for any of the lines, apart from line *At*LORE-OE7-1 (**Fig S1**). *At*LORE-OE7-1 was excluded from further analysis because it exhibited a dwarf, sterile phenotype directly after regeneration from the callus and remained developmentally impaired (**Fig S2, S3**). Although leaf shapes were comparable between all lines, *At*LORE-OE3-4 displayed a slightly sharper serration than the wild type and other *At*LORE-OE or EV lines (**Fig S3**).

**Figure 1.**
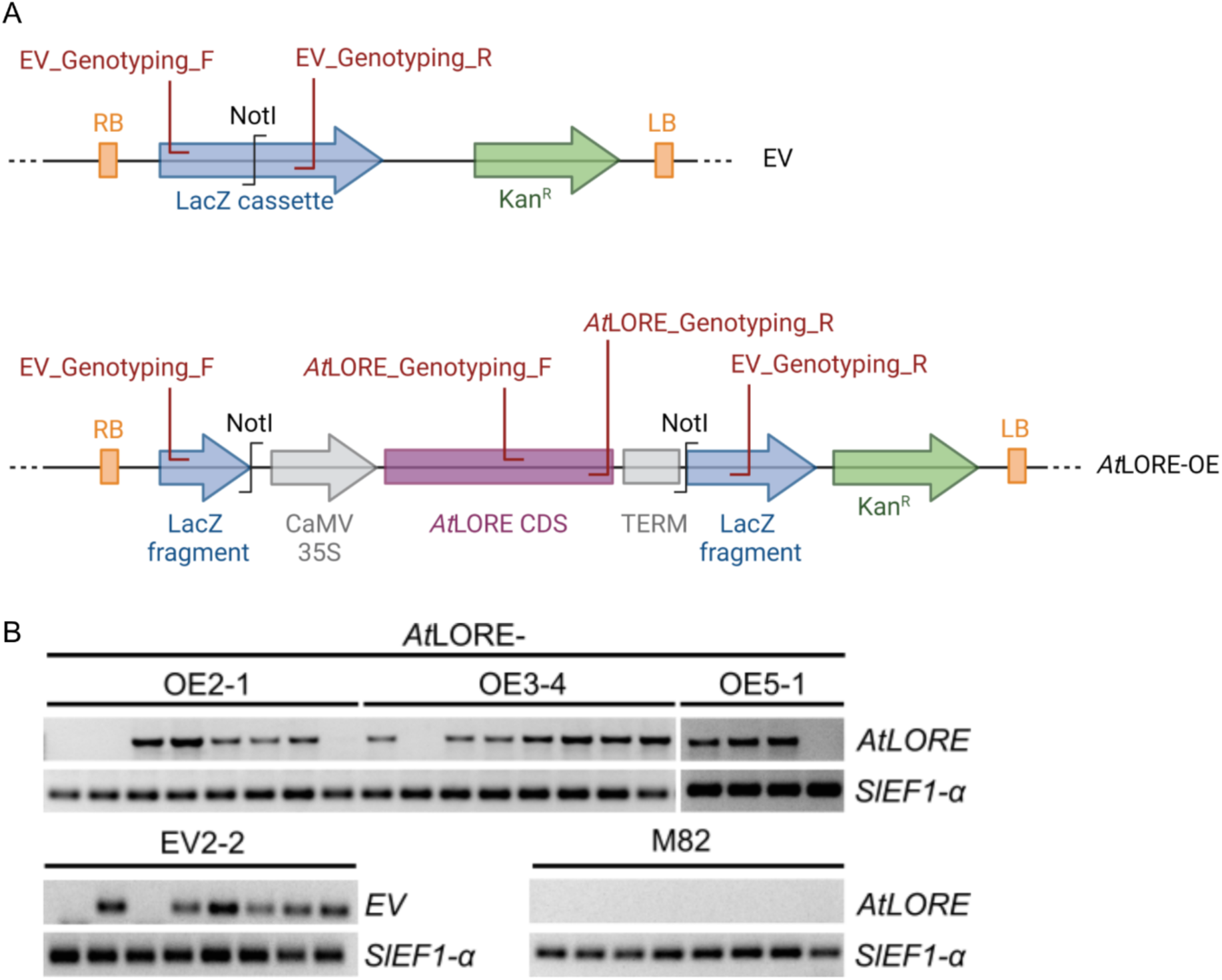
Expression cassettes and genotyping results of transgenic *AtLORE*-OE and EV *S. lycopersicum* lines. **A** Schematic illustration shows the expression cassettes of the empty vector (EV, pART27) and for *At*LORE overexpression (OE), including primers (red) used for genotyping via PCR. *AtLORE* coding sequence (CDS, lilac), cauliflower mosaic virus 35S promotor sequence (CaMV 35S, grey) and an octopine synthase terminator sequence (TERM, grey) were integrated into pART27 via NotI and disrupted the LacZ selection marker cassette (blue). The T-DNA furthermore contains a Kanamycin resistance cassette (Kan^R^, green), and left (LB) and right borders (RB, orange). Scheme was created with BioRender.com. **B** Image shows agarose gel analysis of genotyping PCR from segregating progeny of one EV and three independent *At*LORE-OE lines; untransformed wild type M82 served as control. DNA fragments specific to the T-DNA insertions (*EV* or *AtLORE*) PCR-amplified from extracted gDNA are shown. The wild-type housekeeping gene SlEF1-α was amplified as a control.

For further analysis, the three independent *At*LORE-OE lines and one EV line were grown from harvested seeds. To select transgenic progeny in the segregating T1 generation, we genotyped up to eight seedlings per line by PCR on extracted genomic DNA (gDNA) for the presence of *AtLORE* or EV T-DNA (**Fig 1B**). The housekeeping gene *ELONGATION FACTOR 1 ALPHA* from *S. lycopersicum* (*SlEF1-α*) served as control. For these lines, we evaluated overall macroscopic growth phenotypes (**Fig 2A-C**) and germination rates (**Fig 2D**) to determine whether the transformation process or *AtLORE* overexpression negatively affected their development. All transgenic OE plants grew similarly to M82 wild-type and EV control lines.

**Figure 2.**
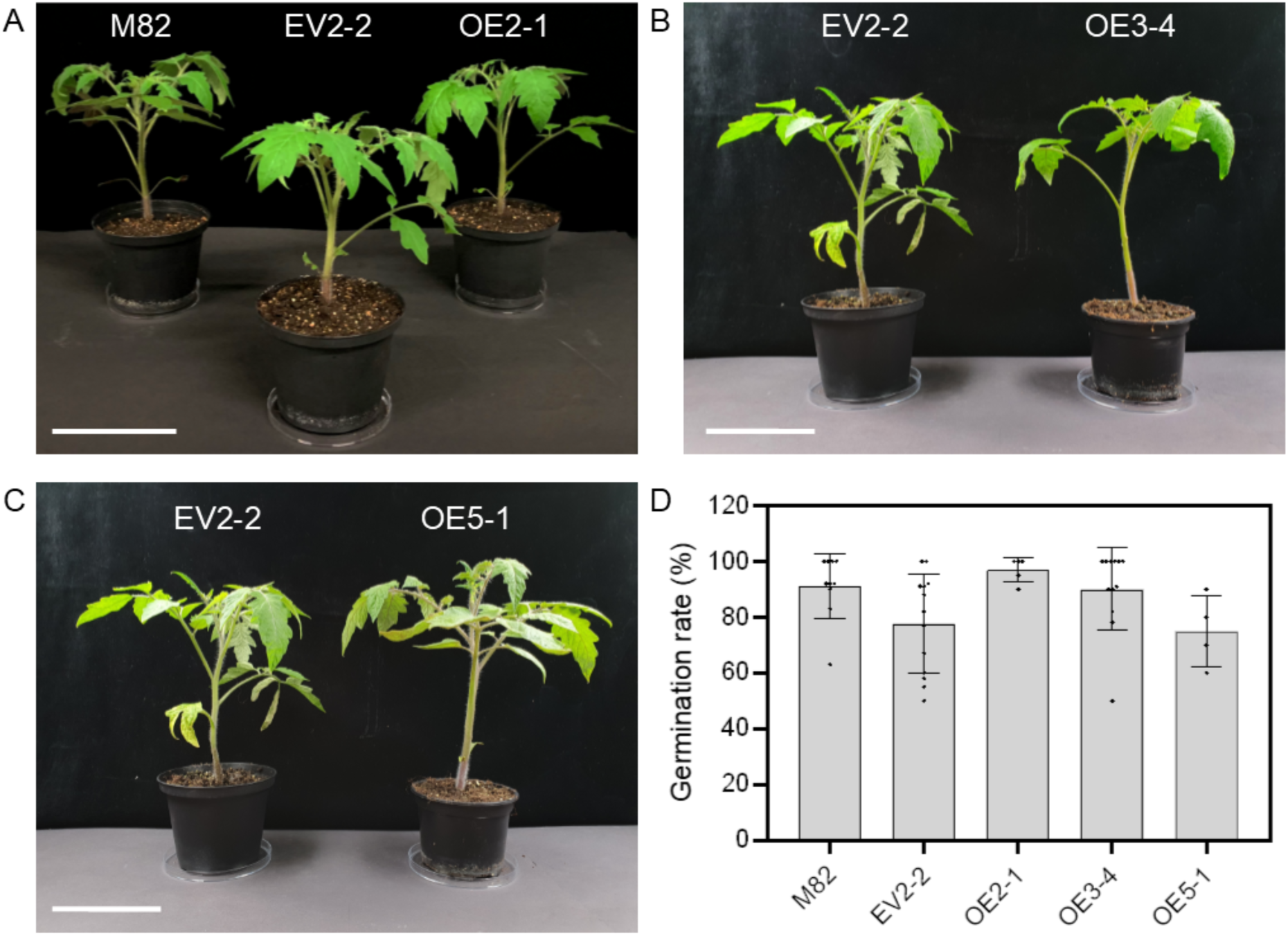
Wild-type and *AtLORE*-transgenic *S. lycopersicum* are phenotypically indistinguishable. **A-C** Photographs of 8-week-old plants of the specified genotypes. Scale bar represents 10 cm. **D** Graph displaying the germination rates (number of germinated seeds/number of seeds sown) of *AtLORE*-transgenic overexpression (OE) and empty vector (EV) lines compared to the M82 wild-type control. Bar graphs show mean with SD of pooled data from at least three biological replicates; each data point corresponds to an individual set of 10 seeds of the indicated line. Germination rates of seeds were assessed one week after sowing. Data do not show significant differences (one-way ANOVA with Tukey’s multiple comparisons test, α = 0.05).

### Overexpression of *AtLORE* in *S. lycopersicum* confers 3-OH-C10:0 sensing

To test whether *AtLORE* overexpression renders tomato sensitive to 3-OH-C10:0, we assessed the production of the phytohormone ethylene as a typical PTI output (Anver & Tsuda, 2015) upon elicitation with 3-OH-C10:0. We analyzed three genotyping positive plants of different *At*LORE-OE lines, EV control or M82 wild type (Fig 3). Leaf discs from *At*LORE-OE3-4 and *At*LORE-OE5-1 plant lines produced high amounts of ethylene when treated with 3-OH-C10:0 compared to the control treatment, while the EV control line showed no detectable ethylene production. For *At*LORE-OE2-1, only two out of three plants responded with rather weak ethylene production compared to *At*LORE-OE3-4 and *At*LORE-OE5-1 (**Fig 3A**). To evaluate the specificity of *At*LORE sensing in *S. lycopersicum,* we tested the highest responding lines *At*LORE-OE3-4#1 and *At*LORE-OE5-1#7 with different concentrations of 3-OH-FAs of different chain lengths. We treated plants with either 1, 5 or 10 µM 3-OH-C10:0, 3-OH-C14:0 or the same volume of EtOH as control. For both lines, we observed a strong concentration-dependent and chain length-specific response (**Fig 3B**), resembling the response characteristics described for *At*LORE in *A. thaliana* (Kutschera *et al*., 2019). This confirms that 3-OH-C10:0 sensing specificity by *At*LORE is transferable from *A. thaliana* to *S. lycopersicum*. Plants OE2-1#17, OE3-4#7 and OE5-1#1, which reacted with high ethylene production, and the control line EV2-2#2 were maintained for further analyses and new stem cuttings were taken from them for further experiments as required.

**Figure 3.**
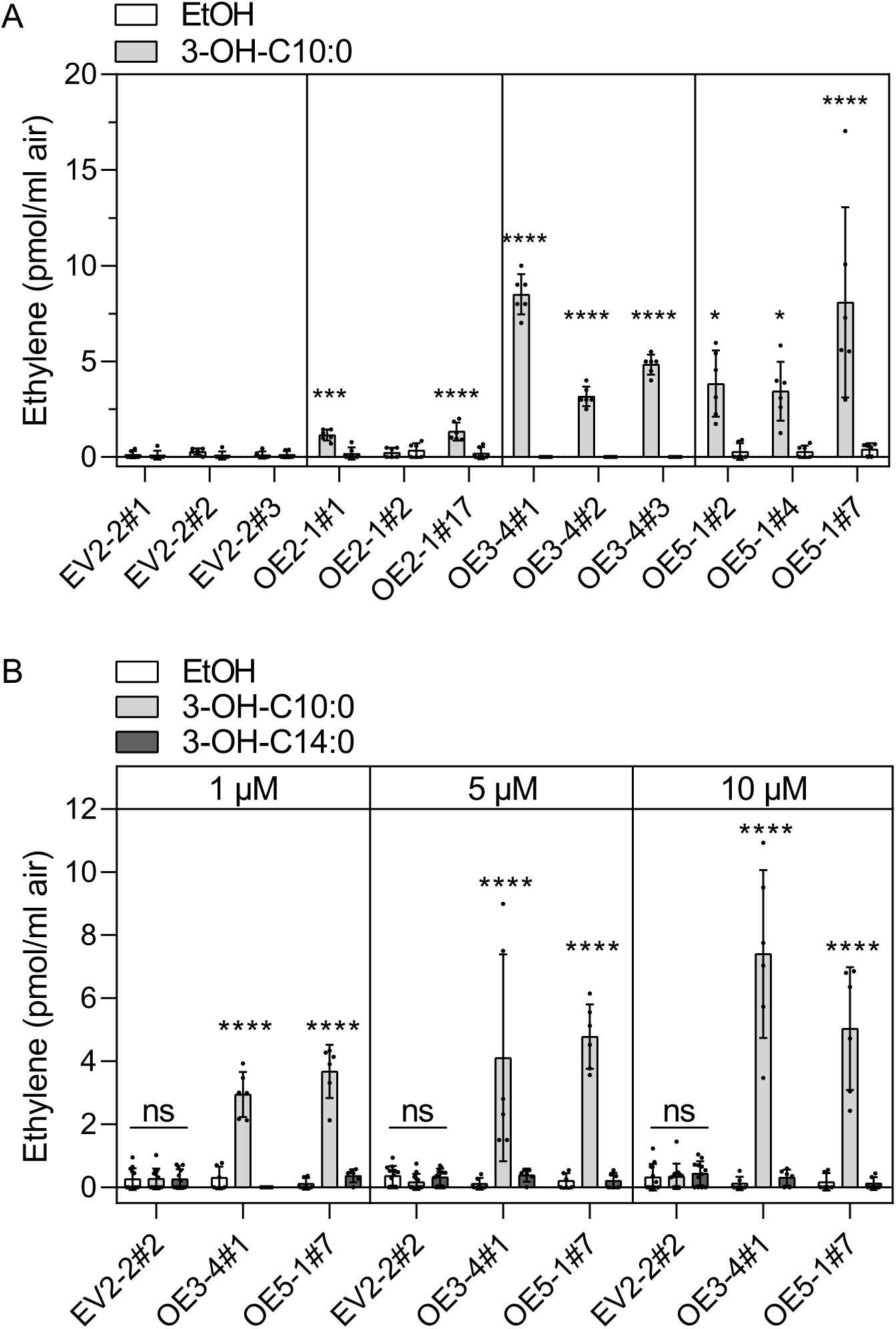
*AtLORE* overexpression in *S. lycopersicum* confers chain length-specific and concentration-dependent 3-OH-FA sensing. **A** Graph displays ethylene production (pmol/mL air) of leaf discs from three independent genotyping-positive plants of each *AtLORE*-transgenic tomato line three hours after elicitation with 5 µM 3-OH-C10:0 or the same volume of EtOH as control. **B** Graph displays ethylene production of the most strongly responding lines from (A), OE3-4 #1 and OE5-1 #7, and an empty vector (EV) control to different concentrations of 3-OH-FAs of different chain lengths. Leaf discs were treated with 1, 5 or 10 µM of 3-OH-C10:0, 3-OH-C14:0 or EtOH as a control for three hours. **A, B** Bar graphs show mean with SD of pooled data from two biological replicates. Individual ethylene measurements per line are represented by black dots (n≥6). Statistical differences between treatments were analyzed by two way-ANOVA with Tukey‘s multiple comparisons test for each line (A) or concentration (B); ****, P<0.0001; ***, P≤0.001; **, P≤0.01; *, P≤0.05; not significant (ns), P≥0.05.

### Overexpression of *AtLORE* in *S. lycopersicum* increases resistance towards pathogenic *P. syringae* pv. *tomato* DC3000

To test whether overexpression of *AtLORE* leads to increased disease resistance in tomato, we performed pathogen infection assays by spray inoculation of *At*LORE-OE and EV plants with *Pst* DC3000. In line with the ethylene accumulation data (Fig 3), bacterial titers were significantly lower in the strongly responding lines OE3-4#1 and OE5-1#7 compared to the EV control line three days after inoculation (**Fig 4A**). These strongly responding lines also showed fewer disease symptoms on leaves (**Fig 4B**) compared to the EV control. No statistically significant reduction in bacterial titers was measurable in the low ethylene-producing line OE2-1#17 three days after infection. However, slightly reduced macroscopic disease symptoms were repeatedly observed in this line compared to the EV control (**Fig 4B**). Our results demonstrate that overexpression of *AtLORE* in tomato might be an effective disease management strategy to increase resistance to bacterial pathogens without compromising plant growth and development.

**Figure 4.**
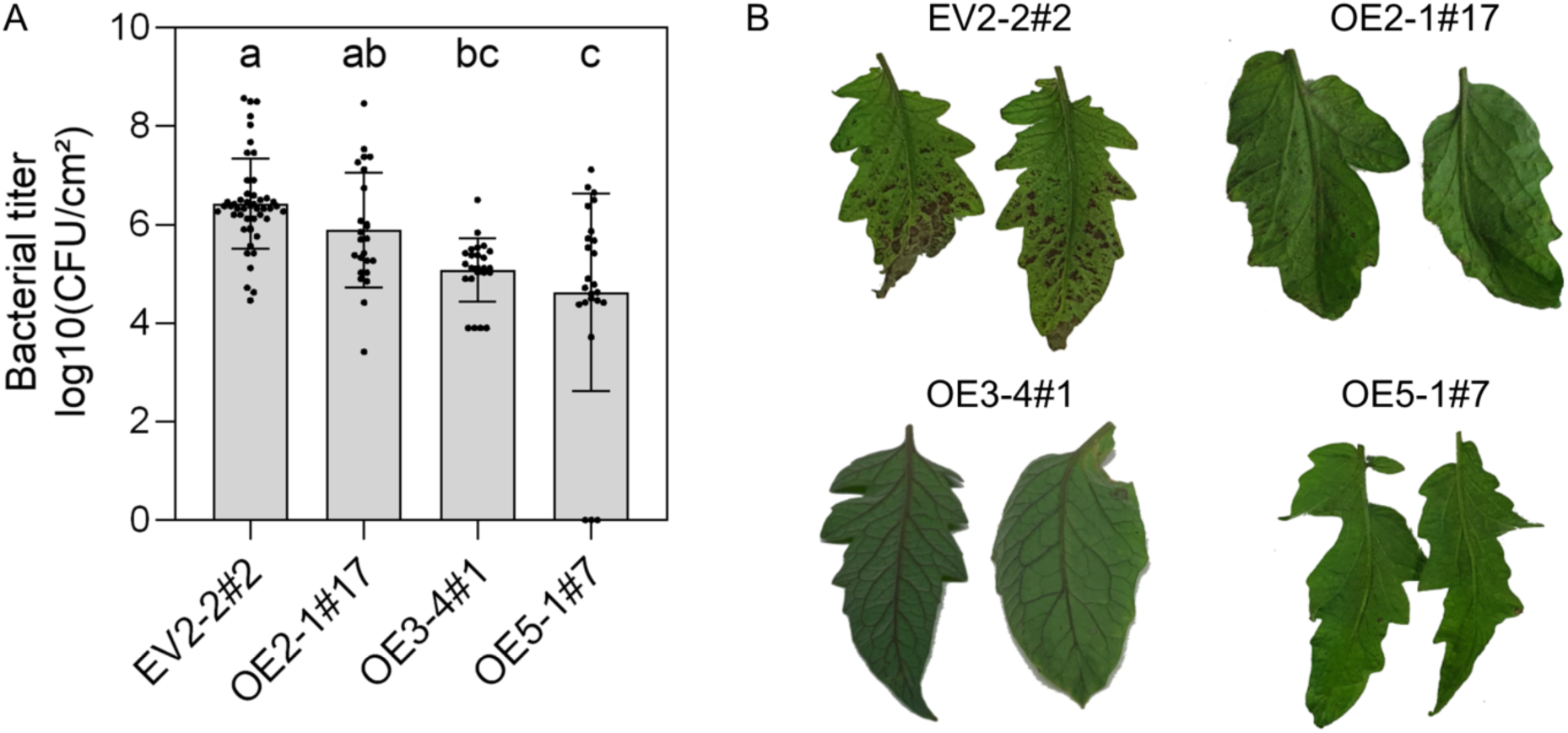
*At*L*ORE* overexpression enhances resistance to *P. syringae* pv. *tomato* (*Pst*) DC3000. *Pst* DC3000 infection of *AtLORE*-transgenic *S. lycopersicum* lines was assessed three days after spraying leaves of the indicated lines with a bacterial suspension of an optical density OD_600_ of 0.002. Experiments were performed on cuttings four to six weeks after their propagation. **A** Graph displays the bacterial titers in leaflets from three cuttings per transgenic line. Bar graphs show mean with SD of pooled data from two independent biological replicates with black dots representing individual data points (n=12 samples from four leaves per cutting and replicate). Statistical differences were analyzed by one-way ANOVA and Tukey’s multiple comparisons test, α=0.01. **B** Photographs show leaflets of the indicated transgenic tomato lines three days after *Pst* DC3000 infection and illustrate the macroscopic disease symptoms.

### Pretreatment with 3-OH-C10:0 activates systemic and broad-spectrum disease resistance

To investigate whether pretreatment of *AtLORE*-transgenic tomato with 3-OH-C10:0 before infection induces systemic resistance against filamentous pathogens, we treated cuttings of EV2-2#2 and OE3-4#1 with 10 µM 3-OH-C10:0 dissolved in water or a water control via soil irrigation. After 48 hours, we spray-inoculated leaves with sporangia solutions of *P. infestans* Pi100 or spore solutions of *A. solani* NL03003 and quantified the infection frequency 14 days post-inoculation (Fig 5). Compared to control-treated plants or the EV control (EV2-2#2), 3-OH-C10:0-treated *At*LORE-OE3-4 plants were significantly more resistant to *P. infestans*. Interestingly, for *A. solani*, we observed reduced infection of *At*LORE-OE3-4 independently of the pretreatment with 3-OH-C10:0 (Fig 5). The infection score of the water-treated *At*LORE-OE3-4 plants was significantly lower than that of the EV control. Pre-treatment of plants with 3-OH-C10:0 further significantly enhanced resistance to *A. solani* compared to the water-treated plants. These data indicate that overexpression of *AtLORE* enables tomato to activate systemic resistance by sensing 3-OH-C10:0, thereby enhancing resistance against filamentous pathogens.

**Figure 5.**
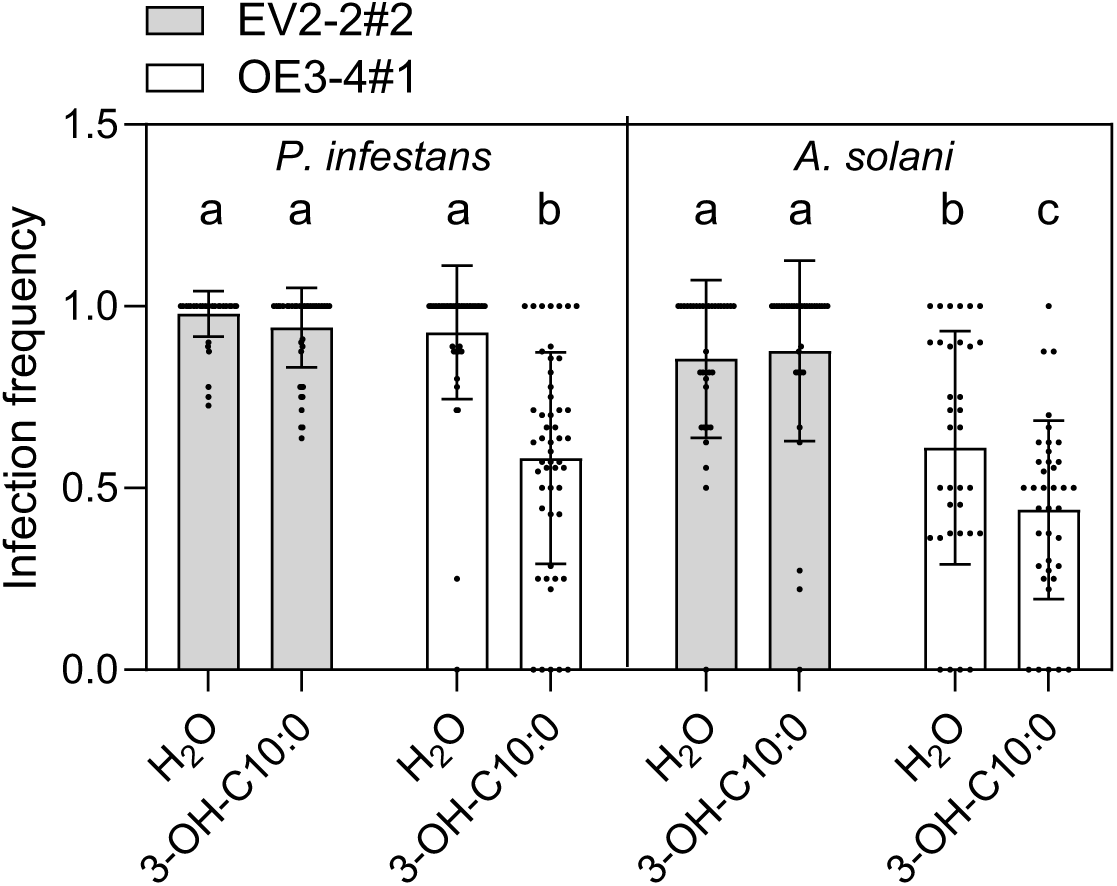
Pretreatment with 3-OH-C10:0 induces resistance against *P. infestans* and *A. solani*. Graph displays the infection frequency (number of symptomatic leaves/infected leaves) 14 days after spray inoculation with *P. infestans* Pi100 or *A. solani* NL03003 of two to three cuttings from EV or *At*LORE-OE3-4, that were pretreated for 48 h via soil irrigation with 10 µM 3-OH-C10:0 dissolved in water or a water-only control. Experiments were performed six to seven weeks after propagation by cutting. Pooled data from two biological replicates are shown, which were obtained from five to six stem cuttings each (n=number of assessed leaves, n≥34 for *A. solani*, n≥50 for *P. infestans*). Statistical differences were analyzed by one-way ANOVA and Tukey’s multiple comparisons test, α=0.05, for *A. solani* and *P. infestans*, respectively.

## DISCUSSION

Historically, resistance breeding has relied primarily on the introduction of major *R* genes from cultivars or landraces of the same species into a variety lacking the desired trait. In recent years, however, several studies have highlighted the potential of cross-family PRR transfer as a promising alternative for engineering disease resistance in crops (Lacombe *et al*., 2010; Mendes *et al*., 2010; Afroz *et al*., 2011; Bouwmeester *et al*., 2014; Tripathi *et al*., 2014; Lu *et al*., 2015; Schoonbeek *et al*., 2015; Schwessinger *et al*., 2015; Hao *et al*., 2016; Wang *et al*., 2016; Boschi *et al*., 2017; Tripathi *et al*., 2017; Kunwar *et al*., 2018; Mitre *et al*., 2021; Piazza *et al*., 2021). These studies have demonstrated that signaling networks downstream of PTI are often sufficiently conserved, even between monocot and dicot plant families, to allow foreign PRRs to fit seamlessly into endogenous signaling pathways of a given species (Afroz *et al*., 2011; Holton *et al*., 2015; Schoonbeek *et al*., 2015; Schwessinger *et al*., 2015; van Esse *et al*., 2020).

Our study shows that the cross-family PRR transfer of the Arabidopsis PRR *At*LORE to tomato enhances resistance against three major tomato diseases, bacterial speck, early blight and late blight, either directly or upon application of synthetic 3-OH-FAs. This suggests that all essential signaling components required for LORE-mediated immunity are present in tomato, although LORE is a Brassicaceae-specific PRR (Ranf *et al*., 2015). Indeed, the signaling components downstream of LORE known to date, *At*PBL34/35/36, *At*RIPK and *At*LOPP, all have putative tomato orthologues (Luo *et al*., 2020; Wang *et al*., 2023). Most of the known PRRs require universal co-receptors for ligand binding and signaling, such as members of the SERK (somatic embryogenesis receptor-like kinase) family, which are usually sufficiently conserved between species, as suggested by the overall functional conservation of SERKs in rice and Arabidopsis for signaling of the PRRs *At*EFR and *Os*Xa21(Chen *et al*., 2014; Holton *et al*., 2015). In contrast, no co-receptors have been described for *At*LORE signaling. Instead, we previously showed that receptor homomerization is essential for 3-OH-C10:0-induced immunity in *A. thaliana* (Eschrig *et al*., 2024). The concept of receptor homomerization could be advantageous to ensure its functionality when transferred into taxonomically different plant families, as co-receptors of phylogenetically restricted receptors may have co-evolved and be absent in other species.

Overexpression or constitutive activation of immunity components is often associated with fitness costs, as plants need to fine-tune the division of their limited resources between immunity and growth (Karasov *et al*., 2017; Wang *et al*., 2021; Zhang *et al*., 2023). Such observations were, for example, made for overexpression of the immune regulator NON EXPRESSOR OF PR1 (NPR1) in rice, which increases disease resistance but also causes growth retardation and spontaneous cell death (Chern *et al*., 2005). Cross-family transfer of PRRs seems to be largely tolerated by recipient plants (Huang & Zimmerli, 2014; Ranf, 2018; van Esse *et al*., 2020). However, for DORN1-transgenic potato, impaired plant and tuber development were reported (Bouwmeester *et al*., 2014). Overexpression of *AtLORE* in tomato does not alter growth, development or reproduction, indicating that LORE expression and signaling are sufficiently controlled in this heterologous system and do not cause autoimmunity or cell death. Interestingly, we previously reported that strong transient overexpression of *AtLORE* in agroinfiltrated *N. benthamiana* leaves leads to receptor auto-activation through homomerization and induces cell death (Eschrig *et al*., 2024). Thus, although *N. benthamiana* and *S. lycopersicum* both belong to the Solanaceae family, our data show that tomato tolerates *AtLORE* overexpression without apparent adverse side effects. This may be due to more moderate expression levels from stably integrated transgenes, or due to the effects of negative regulators that are present in tomato, but absent, diversified or expressed at lower levels in *N. benthamiana*.

We found that 3-OH-C10:0 pretreatment of roots via soil drainage induces resistance against the major tomato pathogens *P. infestans* and *A. solani* in *AtLORE*-transgenic tomato (Fig. 5). Fighting plant diseases through defense priming strategies is a widely discussed disease management approach and offers a great addition to genetic crop protection (Conrath *et al*., 2015; Alexandersson *et al*., 2016; Sandroni *et al*., 2020; Abbasi *et al*., 2021). Primary infection, application of beneficial microbes and treatment with natural or synthetic chemicals activate systemic resistance and switch the plant to a primed state, leading to a more rapid and enhanced resistance activation against subsequent secondary infections (Conrath *et al*., 2015; Abbasi *et al*., 2021; Vlot *et al*., 2021). Similar to 3-OH-C10:0 shown here, other fatty acids, such as eicosapolyenoic fatty acids, have been described as systemic resistance inducers. These fatty acids are released during oomycete infection (or are produced by brown seaweed *Ascophyllum nosodum*) and induce resistance against *Phytophthora capsici* in tomato and pepper (Dye & Bostock, 2021; Lewis *et al*., 2023). Furthermore, soil application of hexanoic acid induces resistance against *Botrytis cinerea* and *P. syringae* in *A. thaliana* and *S. lycopersicum* (Vicedo *et al*., 2009; Kravchuk *et al*., 2011; Scalschi *et al*., 2013). However, for most stimulants, the exact mechanisms of perception and resistance are not understood, so their efficacy in different plant species can hardly be predicted. In contrast, the elicitor, receptor and parts of the immune signaling pathways are characterized for LORE, which could make mc-3-OH-FAs superior to other known resistance inducers. Transgenic expression of *AtLORE* and mc-3-OH-FA soil application could be combined in several important crop species. Yet, the potential effects of mc-3-OH-FAs on plant growth, yield, consumers, and the environment require further evaluation.

Mc-3-OH-FAs, as sensed by LORE, are widespread in nature and can be naturally found in soils, humans, animals, insects, plants, microorganisms and dairy foodstuff (Schildknecht & Koob, 1971; Keinänen *et al*., 2003; Sjögren *et al*., 2003; Jenske & Vetter, 2009; Nagahashi *et al*., 2010; Jones & Bennett, 2011; Kodai *et al*., 2011; Nagahashi & Douds, 2011; Suzuki *et al*., 2013; Mikkelsen *et al*., 2022). Therefore, adverse effects on the environment might be low. With few exceptions, Gram-negative bacteria contain large amounts of 3-OH-FAs as part of lipopolysaccharide (LPS), the main outer membrane component (Alexander & Rietschel, 2001). A common LPS remodeling mechanism is de-acylation, which removes 3-OH-FAs from the lipid A moiety of LPS (Geurtsen *et al*., 2005). Additionally, different bacteria use mc-3-OH-FAs as building blocks in various other compounds, such as lipopeptides (Souza *et al*., 2003; Raaijmakers *et al*., 2006), HAAs and rhamnolipids (Abdel-Mawgoud *et al*., 2010), N-acyl-homoserine lactone-type quorum sensing molecules (Williams, 2007; Thiel *et al*., 2009), or polyhydroxyalkanoates (Raza *et al*., 2018; Paduvari *et al*., 2024). While most of these compounds do not activate LORE-mediated immunity directly (Kutschera *et al*., 2019), mc-3-OH-FAs could be released from these compounds or the ACP/CoA-bound precursors (Zheng, Z. *et al*., 2004; Zheng, Zhong *et al*., 2004; Geurtsen *et al*., 2005; Ernst *et al*., 2006; Kutschera *et al*., 2019; Gerster *et al*., 2022). Free 3-OH-C10:0 FAs are, for example, prevalent in the secretome of Pseudomonas spp. (Schellenberger *et al*., 2021) and the culture medium of *Escherichia coli* (Zheng, Z. *et al*., 2004; Zheng, Zhong *et al*., 2004). Thus, we assume that *At*LORE confers resistance against *P. syringae* upon release of free 3-OH-C10:0 from the pathogen. So far, the release of free mc-3-OH-FAs has not been specifically described for fungi or oomycetes, which are rather known to produce complex, long-chain hydroxy fatty acids (Ivanova *et al*., 2010; Neri *et al*., 2023). However, we found that *A. solani* infection was decreased in *AtLORE*-transgenic tomato independent of the elicitor pretreatment. Thus, *A. solani* potentially directly activates LORE-dependent immunity, possibly by releasing mc-3-OH-FAs or related compounds. So far, only the release of non-hydroxylated medium-chain FAs has been reported in the white-rot fungus *Trametes versicolor* (Hao & Barker, 2022). Medium chain 2- and 3-OH-FAs from plant root exudates affect the growth of arbuscular mycorrhiza fungi (Nagahashi & Douds, 2011) and treatment with LPS (containing 3-OH-C10:0) modulates fungal secondary metabolism (Khalil *et al*., 2014), suggesting that some fungi might sense and respond to mc-3-OH-FAs directly. Furthermore, it has been shown that 3-OH-C10:0 itself has antifungal properties against yeasts and moulds (Sjögren *et al*., 2003). In our study, however, synthetic 3-OH-C10:0 was applied to the soil two days before the leaves were infected with *A. solani* and *P. infestans*. This spatial and temporal separation makes it unlikely that direct contact of 3-OH-C10:0 with the pathogens led to the observed resistance phenotypes. Hence, we assume that the soil application of 3-OH-C10:0 triggers systemic immunity in *AtLORE*-transgenic tomato and subsequently enhances pathogen resistance. Additionally, independent of treatment with synthetic 3-OH-C10:0, the release of mc-3-OH-FAs from soil/plant microbiota or root exudates may lead to low constitutive immune activation in *AtLORE*-transgenic tomato. This might explain or contribute to the observed resistance to *A. solani* and *P. syringae* but might be insufficient for the more aggressive *P. infestans*.

Our study suggests that mc-3-OH-FAs seem to be potent resistance-inducing bio-stimulants on *AtLORE*-expressing plants. In combination, this might form an effective, transferable resistance module and adds to the list of successful examples of resistance engineering in crops. *LORE* might thereby be especially applicable for PRR transfer, as the receptor is not widespread in the plant kingdom compared to others, such as FLS2 (Yue *et al*., 2012), and preexisting natural adaptation of pathogens may be low. This may delay the potential selection of pathogens with altered mc-3-OH-FA profiles that would evade LORE-dependent immunity, potentially leading to more durable resistance of *LORE*-transgenic crops in the field. In the future, infection assays with *Clavibacter michiganensis* subsp. *michiganensis* and *R. solanacearum* on *AtLORE*-transgenic tomato could confirm its effectiveness against other devastating bacterial tomato pathogens. It would be furthermore interesting to transfer *LORE* into other crops that are often threatened by bacterial pathogens, such as potato or rice, which could profit from direct activation of LORE-dependent immunity or the combination with mc-3-OH-FAs application. On the pathogen side, analysis of mc-3-OH-FA contents in fungi and oomycetes would give new insights into whether LORE-dependent immunity is more widely triggered by pathogens from different kingdoms. As advances in plant immunity research have shown that ETI and PTI mutually enhance plant immunity, another benefit of cross-family PRR transfer, alongside the broadened PTI response, could be the potentiation of ETI (Ngou *et al*., 2021; Tian *et al*., 2021; Yuan *et al*., 2021a; Yuan *et al*., 2021b). This suggests that a combination of the *At*LORE expression with QTLs or *R* genes may promote even more profound and durable resistance. Initial evidence for this aspect is provided in potato, where an introgressed QTL from wild potato was combined with an *AtEFR* transgene. The study demonstrated an additive effect of quantitative resistance and heterologous PRR expression over QTL or EFR alone (Boschi *et al*., 2017). In this respect, transgenic approaches and gene-editing tools hold great potential to further engineer durable resistance or circumvent growth-immunity trade-offs, as they offer great flexibility in transgene combinations, tailoring ligand specificity, or additional introduction of positive and negative immune regulators (van Esse *et al*., 2020; Luo *et al*., 2021; Cadiou *et al*., 2023). In conclusion, our study shows that the Brassicaceae-specific PRR *At*LORE is a great model for investigating genetic resistance engineering via cross-family gene transfer. It further emphasizes the advances and importance of basic molecular research in the field of plant immunity and underlines its great potential for application in modern and sustainable agriculture in the future.

## EXPERIMENTAL PROCEDURE

### Molecular cloning

The coding sequence (CDS) of *At*LORE (AT1G61380) was amplified from cDNA before (Ranf *et al*., 2015) and cloned via a binary vector system including pART7 and pART27 (Gleave, 1992) for Agrobacterium-mediated transformation. *At*LORE CDS was amplified via PCR with adapters containing enzymatic restriction sites for integration into the multiple cloning site of the primary cloning vector pART7 via XhoI and EcoRI (Thermo Fischer Scientific, Darmstadt, Germany). The CaMV 35S expression cassette of pART7, including the desired CDS, was transferred into the binary vector pART27 via NotI (Thermo Fischer Scientific, Darmstadt, Germany). The empty vector (EV) pART27 or pART27-*At*LORE were transferred into *Agrobacterium tumefaciens* GV3101 pMP90 for Agrobacterium-mediated transformation of tomato.

### Generation of stable transgenic *S. lycopersicum*

Stable transgenic *S. lycopersicum* cv. M82 containing the empty vector pART27 or pART27-*At*LORE were generated under sterile conditions by Agrobacteria-mediated transformation of cotyledons, callus formation and regeneration of transgenic plants as described (Wittmann *et al*., 2016). Fully regenerated sterile transgenic T0 plants were transferred to soil, genotyped via PCR and grown in the greenhouse until seed set.

### Genotyping by PCR

The presence of T-DNA in transgenic plants was confirmed via extraction of genomic DNA and genotyping by PCR with the REDExtract-N-Amp™ Plant PCR-Kit (Merck, Darmstadt, Germany, Product No. XNAPR-1KT). Primer sequences are listed in Supplementary **Table S1**.

### Growth conditions

After transformation and in vitro callus regeneration, transgenic tomatoes were transferred to potting soil (Einheitserde CL ED73, Patzer Erden, Germany) mixed with vermiculite (1:8). Plants were grown in climate chambers (Fitotron SGC120-H, Weiss Technik, Germany) under long-day conditions (16h light/8h dark) at 23°C and 60% relative humidity and transferred to the greenhouse for seed set. Plants grown from seeds were sterilized, germinated on Jiffy pellets (Jiffy-7, 44 mm; Jiffy Products International AS, Norway) and transferred to potting soil/vermiculite (1:8). Plantlets were grown in climate chambers as described above and propagated for experiments using stem cuttings.

### Seed extraction and sterilization

Fruits of T0 tomato were cut open, and fruit flesh was removed, mixed 1:1 with 3N HCl, and incubated with stirring for 20-30 minutes to remove seed coats. Seeds were washed with H_2_O, neutralized in 10% (m/v) Na_3_PO_4_ for 30 minutes, washed again with H_2_O and dried on filter paper over night. Before planting, all seeds were surface sterilized with 3% sodium hypochlorite for 10 min and washed thrice with H_2_O.

### Assessment of tomato seed germination rates

Surface-sterilized tomato seeds were germinated under sterile conditions on water-soaked filter paper in Petri dishes and incubated in a long-day climate chamber (16h light/8h dark). The germination rates (number of germinated seeds/number of sown seeds) were assessed on the basis of radical and hypocotyl emergence one week after sowing.

### Ethylene measurement

A leaf-disc-based method was performed to evaluate ethylene accumulation in the different plant lines as previously described (Kahlon *et al*., 2023). Leaf discs were sampled with a 4 mm diameter biopsy puncher and incubated overnight at room temperature while floating on H_2_O to allow wounding reactions to decline. Three leaf discs were transferred to 5 ml glass vials containing 300 µl of H_2_O. Elicitors (3-hydroxydecanoic acid, Cat. No. J41087, Manchester Organics, Runcorn, United Kingdom; 3-hydroxytetradecanoic acid, Cat. No. M-1735, Cayman Chemial, Michigan USA; both dissolved in EtOH) or the corresponding volume of EtOH as control were added to the vials, which were immediately sealed with septa (Carl Roth GmbH, Germany). Upon 3 hours of incubation with constant shaking (50 rpm, Polymax 2040, Heidolph Instruments GmbH & Co. KG, Schwabach, Germany), 1 ml of air was sampled with a 1 ml syringe and injected into a gas chromatograph (Varian 3300, Waters^TM^, Germany) equipped with an AlO_3_ column (length 1 m). The detector was set to a temperature of 225°C and the column and injector to 80 °C. The gases H_2_, N_2_ and O_2_ at 0.5 MPa each were used to separate ethylene from the sample. The amount of ethylene was calculated based on the standard calculation developed by Von Kruedener (Von Kruedener *et al*., 1995) using the area under the curve (AUC).

### Infection assays with *Pst* DC3000

*Pst* DC3000 was grown on KB medium with 50 µg/ml Rifampicin at 28°C for two days. Bacteria were scratched from plates and diluted in 10 mM sterile MgCl_2_ and 0.04% Silwet L77 (Kurt Obermeier GmbH & Co. KG, Germany) to an OD600 of 0.002. Spray infection assays were performed on 4- to 6-week-old stem cuttings of *At*LORE-OE and EV lines. Four cuttings per line were prepared, from which three were sprayed with *Pst* DC3000 and one with mock (10 mM MgCl_2_, 0.04% Silwet L-77). Plants were covered with bags for two days to maintain high humidity. Three days post infection, disease symptoms were assessed by photography and bacterial colony counting. For each of the three infected cuttings four leaves were sampled by punching one leaf disc (diameter 4 mm) from the same position of three random leaflets, leading to 12 technical replicates. After adding 100 µl MgCl_2_ (10 mM) and two glass beads per sample, all samples were ground for 2 minutes at 25 Hz (TissueLyser II, QIAGEN, Hilden, Germany). A dilution series was performed on this mixture with MgCl_2_ (10 mM) to a dilution of up to 1×10^-8^. 10 µl of each dilution was drop-inoculated on KB-Agar plates (10 µg/ml Rifampicin) and incubated at 28°C for about 36 hours. Colony-forming units were counted, normalized to the leaf area and log10 transformed.

### Induced resistance infection assay with *A. solani* and *P. infestans*

To induce resistance, six- to seven-week-old stem cuttings of EV and *At*LORE-OE lines were irrigated with 10 µM 3-OH-C10:0 dissolved in water (Cat. No. J41087, Manchester Organics, Runcorn, United Kingdom) or a water control 48 hours prior infection. The whole leaf area was spray infected with spore solutions of *P. infestans* isolate Pi100 (3000 sporangia/ml) as described (Kahlon *et al*., 2021) or *A. solani* NL03003 (5000 spores/ml). Leaf infection frequencies (symptomatic leaflets/inoculated leaflets) were assessed 14 days after infection as described (Kahlon *et al*., 2021).

## ACKNOWLEDGEMENTS

We thank Bert Evenhuis for kindly providing the *A. solani* isolate NL03003 and Remco Stam for providing the *P. infestans* isolate Pi100. We thank Kai Steinmetz, Sabine Zuber and Bärbel Breulmann from the TUM Plant Technology Center for maintenance of tomatoes in the greenhouse. Research was supported by grants from the German Research Foundation to S.R. (SFB924/TP B10 and Emmy Noether program RA-2541/1).

## AUTHOR CONTRIBUTIONS

Conception of the project: SR, PK, RH; generation of transgenic tomato: AH, CA; experimental work, data collection, analysis and representation: SE, PK; data interpretation and discussion of results: SE, PK, SR; drafting the manuscript: SE; critical revision of the manuscript: PK, SR, RH, CS.

## DATA AVAILIBILITY STATEMENT

All data supporting the findings of this study are available within the article and its supplementary material. Raw data are available from the corresponding author on request.

## SUPPORTING INFORMATION LEGENDS

**Figure S1 Propagation of *At*LORE-transgenic *S. lycopersicum* cv. M82 in the greenhouse.** Transgenic tomatoes overexpressing *At*LORE or harboring an empty vector control were regenerated from callus culture and grown in the greenhouse (**A**). Most transgenic lines did not show obvious growth, fruiting or yield alterations (**B**, fruits of OE2-1).

**Figure S2 *At*LORE-transgenic tomato line OE7-1 exhibits an impaired growth phenotype.** Line OE7-1 shows a dwarf, developmentally impaired phenotype (**A-B**). The flowers of OE7-1 appeared to be sterile due to an exerted style that outgrows the stigma from the anthers and prevents self-pollination (**C-D**).

**Figure S3 Leaf shape phenotypes of *At*LORE-transgenic tomato lines (T0). Table S1 Primer sequences for genotyping PCR**

