## Supplementary Material for "Cross-family transfer of the Arabidopsis cell-surface immune receptor LORE to tomato confers sensing of 3-hydroxylated fatty acids and enhanced disease resistance"

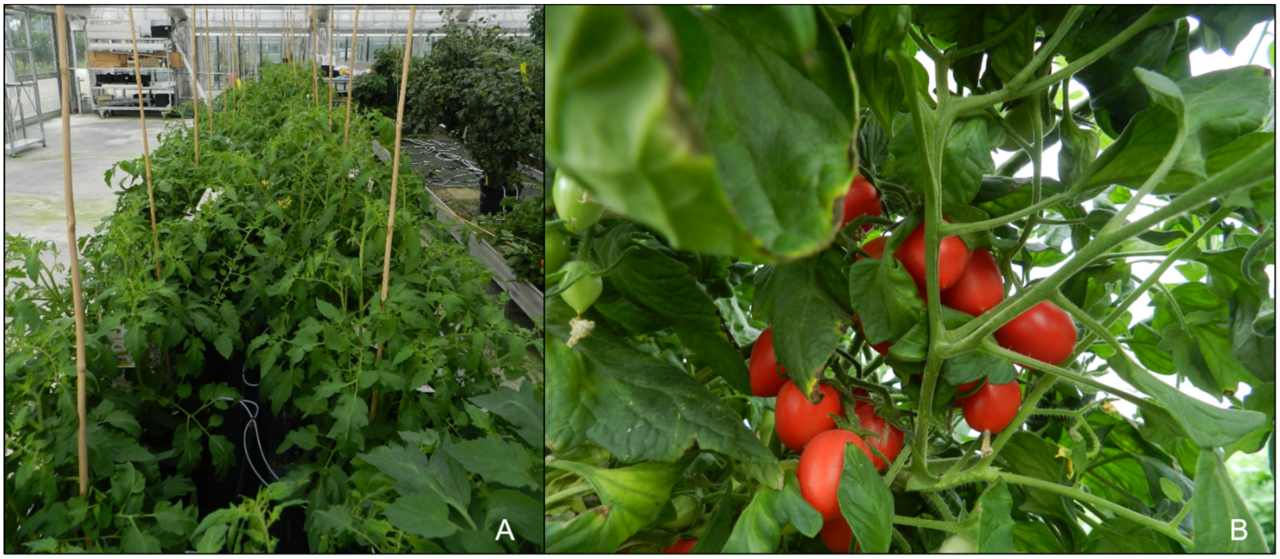

**Figure S1 Propagation of *At*LORE-transgenic *S. lycopersicum* cv. M82 in the greenhouse.** Transgenic tomatoes overexpressing *At*LORE or harboring an empty vector control were regenerated from callus culture and grown in the greenhouse (**A**). Most transgenic lines did not show obvious growth, fruiting or yield alterations (**B**, fruits of OE2-1).

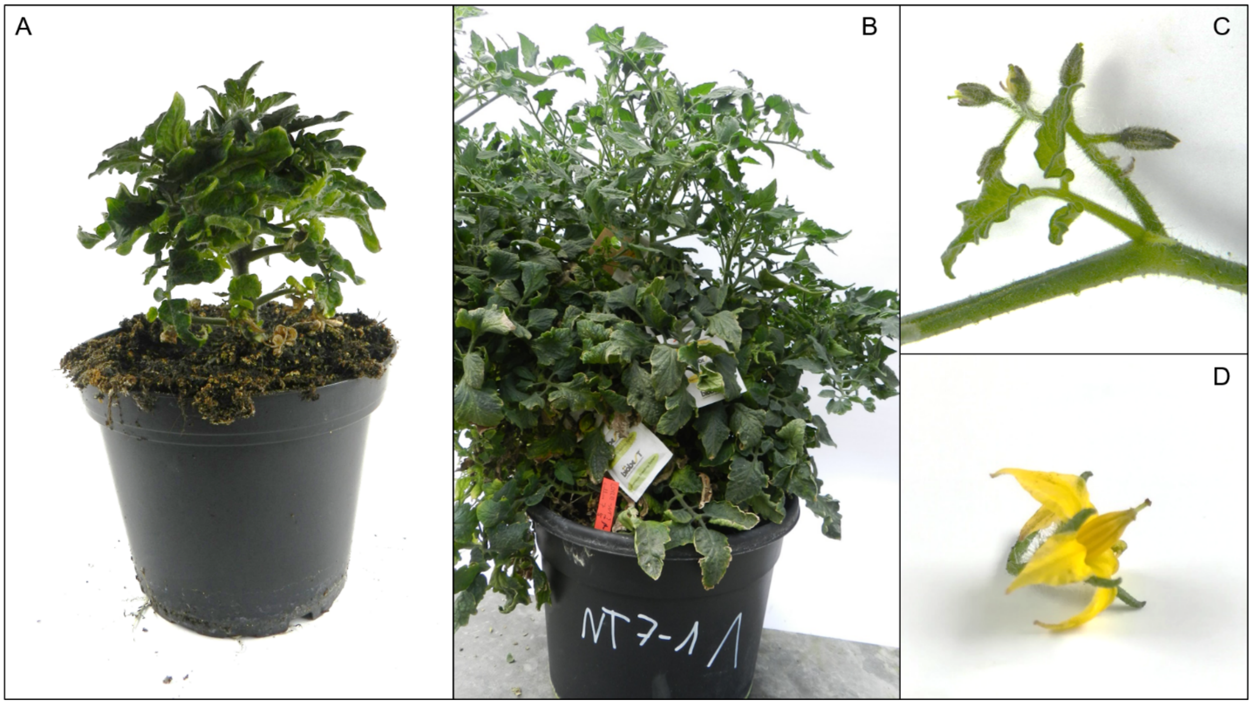

**Figure S2 *At*LORE-transgenic tomato line OE7-1 exhibits an impaired growth phenotype.** Line OE7-1 shows a dwarf, developmentally impaired phenotype (**A-B**). The flowers of OE7-1 appeared to be sterile due to an exerted style that outgrows the stigma from the anthers and prevents self-pollination (**C-D**).

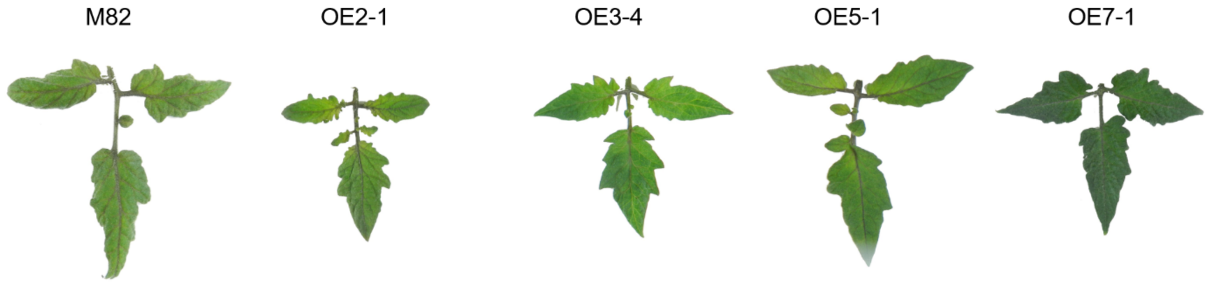


**Figure S3 Leaf shape phenotypes of *At*LORE-transgenic tomato lines (T0).**

**Table S1 Primer sequences for genotyping PCR**

| **Target** | **Primer name** | **Sequence** |
|---|---|---|
| *At*LORE | AtLORE_Genotyping_R | GTACCGAATTCCTACCTTCC |
| | AtLORE_Genotyping_F | GTTCCGGTCAGGGTACAGAG |
| Empty vector T-DNA | EV_Genotyping_F | AACAGCTATGACCATGA |
| | EV_Genotyping_R | GTTTTCCCAGTCACGAC |
| EF1-α | Ef1α_F | TACTGGTGGTTTTGAAGCTG |
| | Ef1α_R | AACTTCCTTCACGATTTCATCATA |
